# Regional pulmonary perfusion, blood volume, and their relationship change during early ARDS in an experimental study

**DOI:** 10.1101/2023.06.19.545593

**Authors:** Arnoldo Santos, Gabriel C. Motta-Ribeiro, Nicolas De Prost, Mauro R. Tucci, Tyler J. Wellman, Marcos F. Vidal Melo, Tilo Winkler

**Author notes:** **Corresponding author** Tilo Winkler, Department of Anesthesia, Critical Care and Pain Medicine, Massachusetts General Hospital and Harvard Medical School, Boston, Mass, USA. Funding: NIH-NHLBI grant R01-HL121228. GCMR was also funded by CAPES, Ministério da Educação do Brasil (scholarship 6344/15-1).

## Abstract

Regional pulmonary perfusion (Q) has been investigated using blood volume (F_b_) imaging as an easier-to-measure surrogate. However, it is unclear if changing pulmonary conditions could affect their relationship. We hypothesized that vascular changes in early acute respiratory distress syndrome (ARDS) affect Q and F_b_ differently. Five sheep were anesthetized and received protective mechanical ventilation for 20 hours while endotoxin was continuously infused. Using dynamic ^18^F-FDG and ^13^NN Positron Emission Tomography (PET), regional F_b_ and Q were analysed in 30 regions of interest (ROIs) and normalized by tissue content (F_bn_ and Q_n_, respectively). After 20 hours, the animals’ lung injury showed characteristics of early ARDS, including gas exchange and lung mechanics. PET images of F_bn_ and Q_n_ showed substantial differences between baseline and lung injury. Lung injury caused a significant change in the F_bn_-Q_n_ relationship compared to baseline (p<0.001). The best models at baseline and lung injury were F_bn_=0.32+0.690Q_n_ and F_bn_=1.684Q_n_–0.538Q_n_^2^, respectively. Early ARDS changed the relationship between F_b_ and Q from linear to curvilinear. Effects of endotoxin exposure on the vasoactive blood flow regulation were most likely the key factor for this change limiting the quantitative accuracy of F_b_ imaging as a surrogate for regional Q.

## Introduction

In healthy lungs, pulmonary perfusion and intravascular blood volume show a vertical gradient caused by gravitation with higher perfusion and blood volume in dependent regions (1,2). However, it is unknown if regional perfusion and regional blood volume have a tight constant relationship or if certain conditions such as endotoxin exposure during early acute respiratory distress syndrome (ARDS) may change their relationship. Clinically relevant would be for example decreases in the regional blood volume resulting in regional vascular collapse causing alveolar dead space and potentially vascular injury, micro-clotting of the blood, or redistribution of perfusion resulting in a mismatch between ventilation and perfusion impairment of pulmonary gas exchange.

During ARDS, the evaluation of pulmonary perfusion is particularly relevant (3) because changes in regional perfusion resulting in a mismatch between ventilation and perfusion worsen gas exchange leading to severe hypoxemia, one of the landmarks of ARDS. During the early phase of ARDS, alterations in the regional conditions including inflammation, vasoconstriction or -dilation, blood clotting, alveolar overdistension, and the intravascular-to-alveolar pressure difference can change the perfusion distribution in the lungs. Besides the effect on gas exchange, changes in pulmonary vascular resistance (PVR) during ARDS can have a clinically significant effect on right ventricular (RV) failure that has been demonstrated to affect patient outcomes (4).

Dual-energy computed tomography (DECT) allows imaging of perfused blood volume, which has been shown to correlate with regional perfusion (5,6) and its availability and speed may have advantages compared to pulmonary SPECT, PET, and CT perfusion imaging methods, e.g. (5,7–12). However, it remains unclear if the relationship between regional blood volume and perfusion could change. For example, it has been shown that hypoxic pulmonary vasoconstriction (HPV) is severely blunted in acute lung injury (11), and regional changes in vasoactive blood flow regulation could affect the relationship between blood volume and perfusion. Also, a PET imaging study reported a curvilinear relationship between blood volume and perfusion in humans (12).

In this study, we hypothesized that lung injury representative of early ARDS with endotoxin exposure is associated with changes in the regional vasoactive blood flow regulation affecting the relationship between blood volume and perfusion. We aimed to study the effects of lung injury on regional perfusion and blood volume in an animal model using PET imaging.

## Material and methods

We performed a detailed new analysis of the regional pulmonary perfusion and blood volume, and changes in their relationship using the original PET imaging data of a previously published experimental study in 6 sheep (13). Here, 5 animals with complete imaging data were included. The Subcommittee on Research Animal Care at the Massachusetts General Hospital approved the experimental protocols.

### Subjects and Experimental design

The experimental design has been previously described in detail (13). Briefly, sheep were anesthetized, intubated and mechanically ventilated using volume control, PEEP 5cmH_2_O, tidal volume 6ml/kg, inspired O_2_ fraction to maintain an arterial oxygen saturation ≥ 90% and respiratory rate to keep a PaCO_2_ between 32 and 45 mmHg.

Further details are described in the online supplement.

Once animal instrumentation was completed, baseline measurements of physiological parameters were obtained. Subsequently, a set of dynamic ^13^NN and ^18^F-FDG PET images was acquired.

After baseline measurements, animals were subjected to a continuous lipopolysaccharide (*Escherichia coli* O5:55, List Biologic Laboratories Inc, USA) infusion of 10ng/kg/min for 20 hours. Then measurements were repeated for lung injury. During the time of the protocol PEEP and FiO_2_ were managed according to the ARDS Network sliding table (low PEEP high FiO_2_)(14).

### PET image acquisition

The PET imaging equipment, protocol, and processing methods have been previously presented in detail (10,15–17), and are described in the online supplement. Briefly, we collected 15 PET transverse slices of 6.5-mm thickness, estimated to encompass approximately 70% of the total sheep lung volume. Reconstructed images consisted of an interpolated matrix of 128 × 128 × 15 voxels with a size of 2.0 × 2.0 × 6.5 mm each. Three different types of PET images were acquired.

1. Transmission scans were obtained at baseline (0 h) and at 20 h using a rotating pin source of ^68^Ge for 10 min. Transmission scans were used for the attenuation correction of the corresponding emission scans, to delineate the lung field, and to determine the fraction of gas (F_gas_).
2. ^13^NN (nitrogen) emission scans were obtained at baseline and at the end of the 20-h mechanical ventilation period for the assessment of regional perfusion (Q_r_), including shunt using the ^13^NN-saline method.
3. ^18^F-FDG emission scans: after ^13^NN clearance, ^18^F-FDG dissolved in 8 ml saline (approximately 40 MBq at 0 h and 200 MBq at 20 h) was infused at a constant rate through the jugular catheter for 60 s.

Further details are provided in the online supplement.

### Image analysis

Lung masks were created from transmission and perfusion scans by thresholding and then manually corrected to assure the adequate selection of lungs. The resulting lung masks were divided in the vertical direction using 15 isogravitational planes and in the axial direction using two sections so that we obtained 30 regions of interest (ROI).

The ROI blood volume was determined by fitting the ^18^F-FDG kinetics of each individual ROI to the three-compartment Sokoloff model (18)

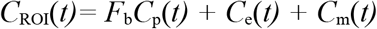

where C_ROI_(t) represents the ROI’s ^18^F-FDG activity in the PET image, F_b_ the blood fraction, C_p_(t) the concentration of FDG in blood plasma, C_e_(t) the concentrations in the extravascular compartment serving as substrate pool for hexokinase, and C_m_(t) the concentration of phosphorylated FDG (18,19).

The normalized tissue fraction (F_tis,n_) of each ROI was first calculated as tissue fraction of the individual ROIs using

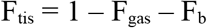

and then normalized by the mean F_tis_ among the ROIs of each individual using F_tis,n_ = F_tis_ / mean(F_tis_).

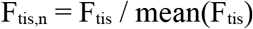

The normalized regional perfusion Q_n_ of each ROI was first calculated as regional perfusion Q_r_ equal to the plateau of the ^13^NN activity reached at the end of the breathhold plus the shunt fraction equal to the relative height of a peak prior to the plateau, and then normalized by the mean perfusion among the ROIs and F_tis,n_

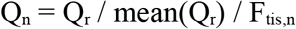

So, regional changes in Q_n_ account for changes in regional lung tissue density. The normalized blood fraction (F_bn_) is based on the blood fraction estimates of the individual ROIs (F_b_) and normalized by the mean blood fraction among the ROIs and F_tis,n_

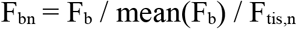

equivalent to the normalization of Q_n_.

To investigate longitudinal changes in regional perfusion and blood volume accounting for changes in cardiac output (CO) and pulmonary blood volume between the lung injury (INJ) and baseline (BL) conditions, adjusted Q_n_ (Q_a_) and F_bn_ (F_ba_) referenced to baseline were calculated using for the lung injury of early ARDS

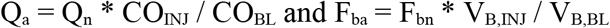

where V_B_ is the overall pulmonary blood volume at lung injury and baseline equal to the sum of the ROIs’ products of blood fraction (F_b_) and mask volume.

For baseline, the adjusted values referenced to baseline are

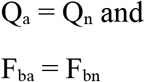

The obtained data were analysed aiming to:

1. Evaluate the effect of the lung injury on the distribution of F_bn_ and Q_n_.
2. Evaluate the relationship between F_bn_ and Q_n_.
3. To test the hypothesis that lung injury modifies the relationship between F_bn_ and Q_n_.
4. To compare the changes in F_bn_ and Q_n_ between baseline and lung injury.

### Statistical analysis

The primary objective of this study was to determine if the relationship between F_b_ and Q during lung injury is different from baseline. To answer it, we performed a kernel density estimator (KDE) test under the null hypothesis that the F_bn_ vs Q_n_ data at lung injury are from the same distribution as the baseline data. As secondary analysis aiming to identify models that describe the relationship between F_bn_ and Q_n_, we compared six different relationships between F_bn_ and Q_n_ at baseline and lung injury conditions and the assignment of F_bn_ and Q_n_ to the independent and dependent parameter using the Bayesian information criterion (BIC) as primary parameter for the identifying the best model. Additional details are described in the online supplement.

A p-value < 0.05 was assumed as significant. Values are expressed as mean ± standard deviation unless otherwise specified. The effect of the early ARDS model in physiologic variables was evaluated by applying a paired t test. The computational analysis was performed using Matlab (Mathworks, Natick, MA). The statistical analysis was performed using Stata (v14.2; StataCorp LLC, College Station, TX), the R Statistical Software (v4.2.1; R Core Team 2022), and the ks package (v1.13.5) (20) for the KDE test.

## Results

### Effects of lung injury representative of early ARDS

After 20 hours of mechanical ventilation and continuous endotoxin infusion the animals included in this analysis showed characteristics of lung injury including deterioration in lung mechanics and gas exchange and also an increase in pulmonary vascular resistance (PVR) (13) (Table 1) (one animal was excluded from the hemodynamic analysis due to a problem with cardiac output measurement, this also applies for adjustment of Q_n_ to baseline cardiac output analysis below).

**Table 1.**
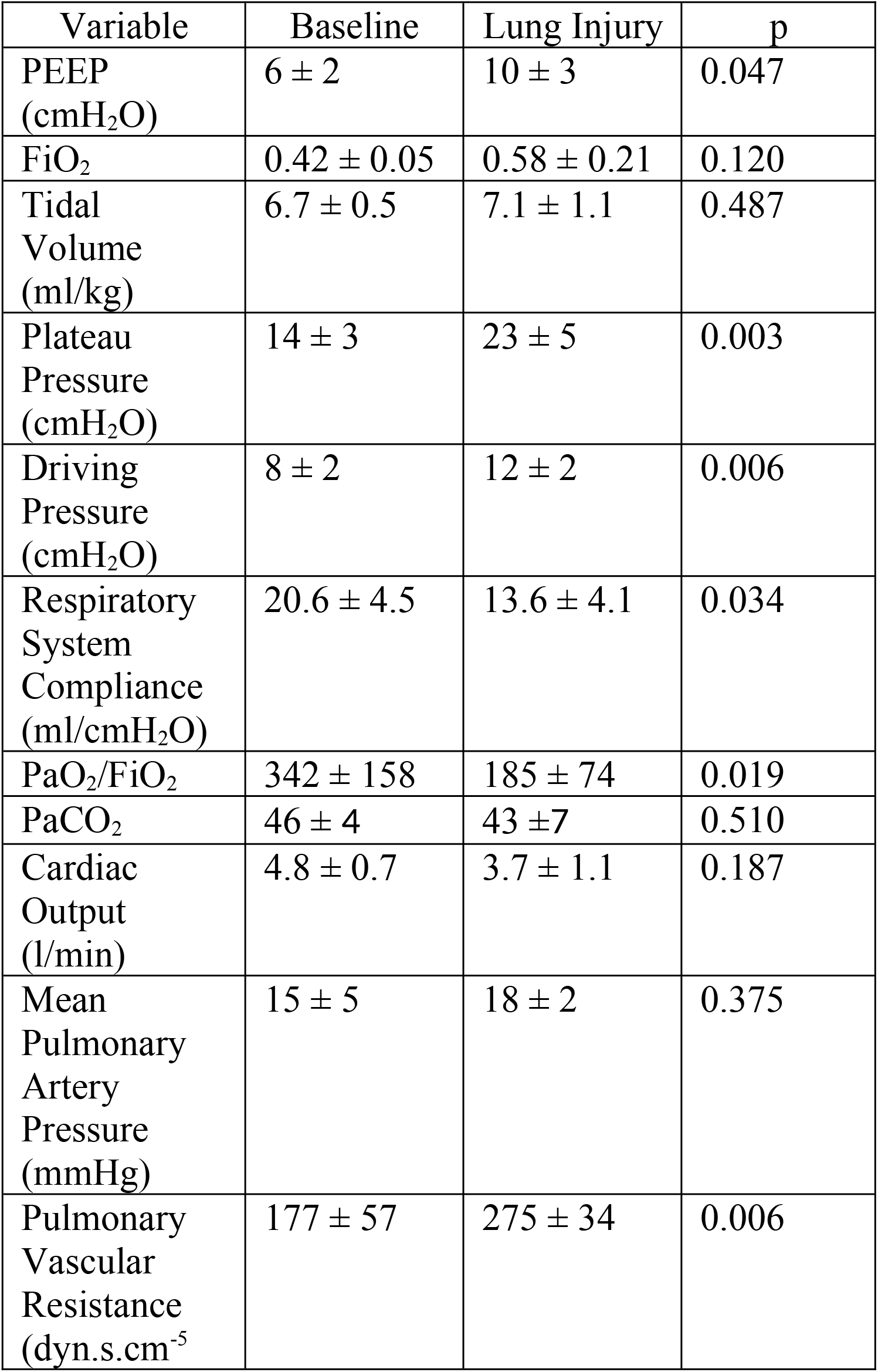
Physiologic parameters and significance tests of differences

PET images of regional perfusion and blood volume show substantial differences between baseline and lung injury (Fig. 1). The vertical distribution of Q_n_ among the ROIs shows at baseline a gradual change from dependent (dorsal) to non-dependent (ventral) regions, but it changes during injury to higher perfusion in dependent and less in non-dependent regions compared to baseline (Fig. 1). The vertical distributions of F_bn_ showed a similar distribution.

**Figure 1.**
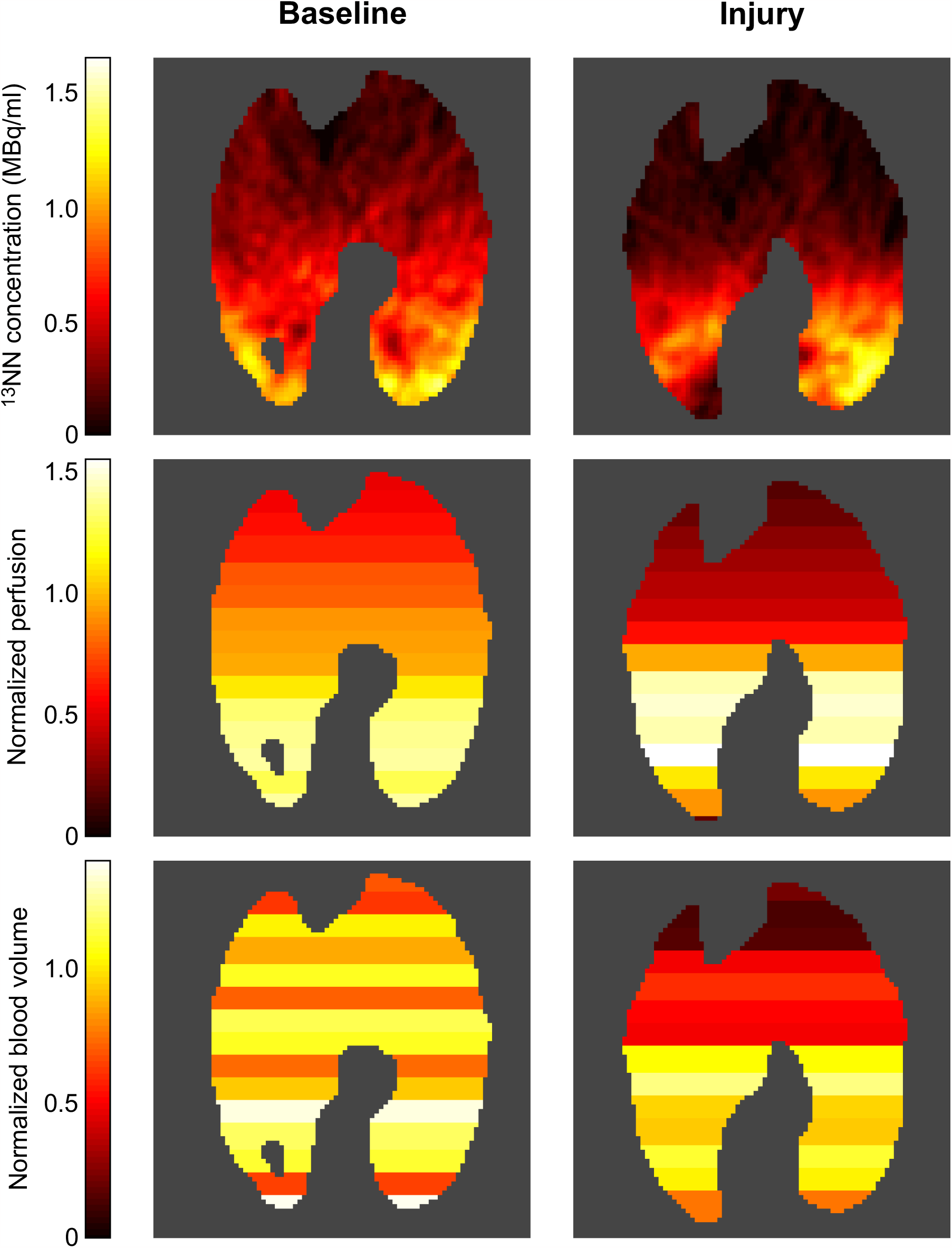
Typical slices of ^13^NN PET images at baseline and lung injury. The ^13^NN concentration in these images is proportional to the regional perfusion of aerated voxels showing a gradual change along the vertical gradient at baseline. In contrast, lung injury resulted in increased perfusion in the dependent regions and very low perfusion in the non-dependent regions. The normalized perfusion of 15 vertically stacked ROIs shows regional differences taking differences in tissue density into account. The distributions of normalized blood volume correlate with the distributions of normalized perfusion. But the images also illustrate the residual variability of the correlation.

### The relationship between F_bn_ and Q_n_ during lung injury representative of early ARDS is different from baseline

Performing a KDE test for the F_bn_-Q_n_ relationship during lung injury compared to baseline, we found a highly significant difference (p<0.001) showing that the F_bn_-Q_n_ point clouds at the two time points are not from the same distribution. After the global test had established this difference, we explored the functional relationships using different mathematical models.

### Characterization of the relationships between F_bn_ and Q_n_

Using the BIC to compare different models, we identified the linear relationship F_bn_ = 0.32 +0.690Q_n_ as the best model for baseline (Fig. 2A, Table 2A). In contrast, the quadratic relationship F_bn_ = 1.684Q_n_ - 0.538Q _n_ ^2^ was the best model during lung injury (Fig. 2B, Table 2B). Additionally, a likelihood ratio test comparing linear and quadratic models resulted in chi-square of 3.76 (p = 0.0526) for baseline and 76.91 (p < 0.001) for lung injury, reinforcing the relevance of quadratic model at early ARDS. The criteria of lowest BIC also suggested better model fitting using F_b_ as dependent and Q as independent variables than the opposite configuration (online supplement, Tables 2A and 2B). The residual scattering was higher at baseline than lung injury.

**Figure 2.**
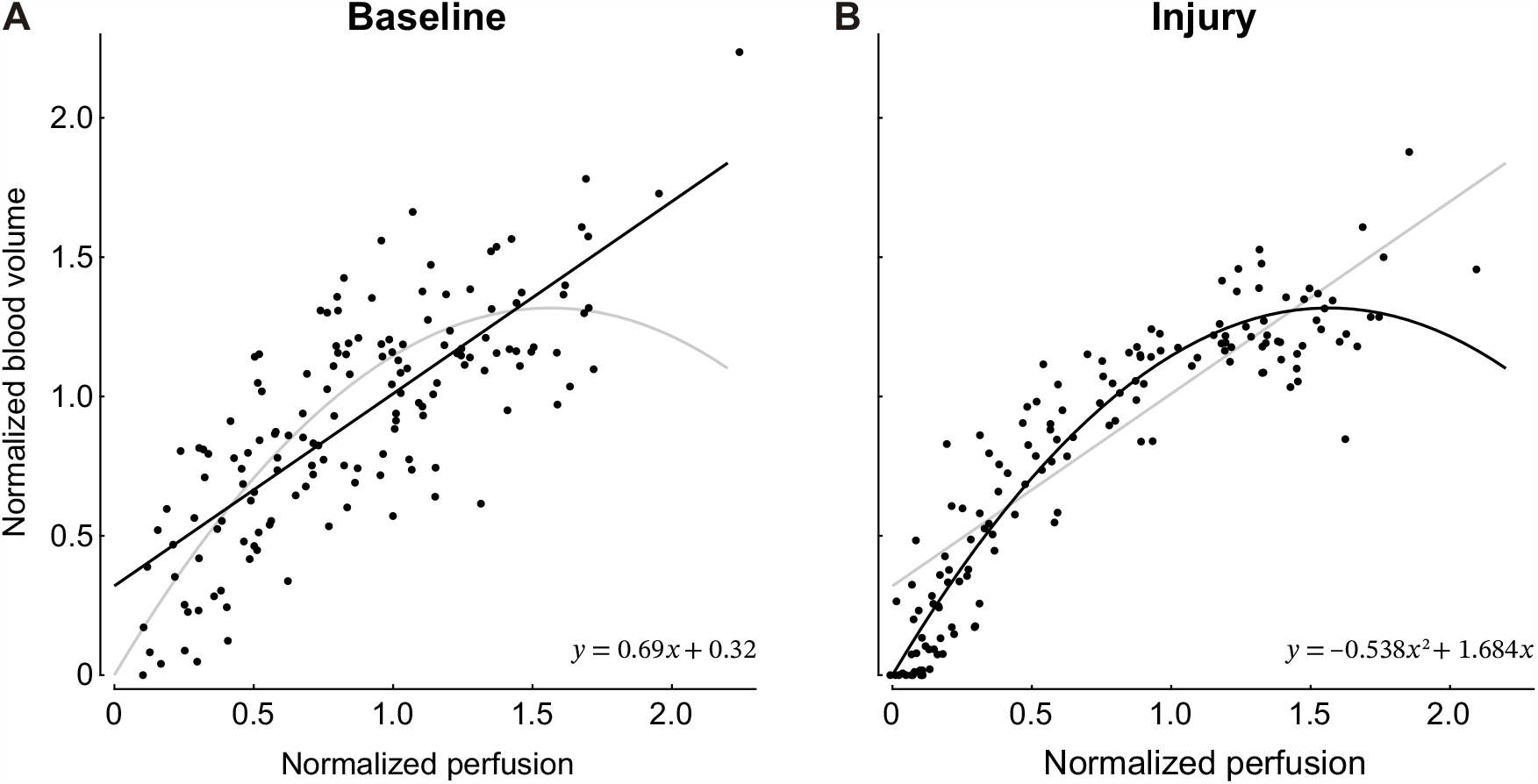
Blood volume vs. perfusion of each ROI at baseline (A) and lung injury (B) show the difference between the two conditions, which a KDE test confirmed as highly significant. Model identification resulted in the selection of a linear relationship as the best model (solid line) for baseline (A) and a curvilinear quadratic relationship for lung injury (B). Each panel also includes the fitted model of the other condition (grey lines) for reference. Note the substantially higher scattering of the data points at baseline compared to lung injury.

**Table 2.** Model comparisons A – Baseline

| Model | a | b | c | BIC |
| --- | --- | --- | --- | --- |
| $y = ax$ | 0.980 | | | 62.381 |
| $y = a + bx$ | 0.320 | 0.690 | | 23.977 |
| $y = ax + bx^2$ | 1.433 | -0.357 | | 27.596 |
| $y = a + bx + cx^2$ | 0.204 | 1.008 | -0.168 | 25.231 |
| $y = e^{ax}$ | 0.078 | | | 156.465 |
| $y = ae^{bx}$ | 0.503 | 0.650 | | 38.425 |

B – Lung Injury
| Model | a | b | c | BIC |
| --- | --- | --- | --- | --- |
| $y = ax$ | 0.967 | | | 10.437 |
| $y = a + bx$ | 0.214 | 0.780 | | -30.598 |
| $y = ax + bx^2$ | 1.684 | -0.538 | | -106.026 |
| $y = a + bx + cx^2$ | 0.004 | 1.674 | -0.532 | -101.042 |
| $y = e^{ax}$ | 0.097 | | | 228.320 |
| $y = ae^{bx}$ | 0.415 | 0.775 | | 38.902 |

### Regional changes in Q_n_ and F_bn_

In nondependent zones, where Q_n_ was already low at baseline, the lung injury caused consistently further decreases in perfusion (Fig. 3A). In these zones, the changes in F_bn_ were less consistent, but showed predominantly lower F_b_ during lung injury than baseline (Fig. 3B). In dependent zones, F_bn_ and Q_n_ were in most cases higher during injury than baseline, but this effect was more consistent and marked for Q_n_.

**Figure 3.**
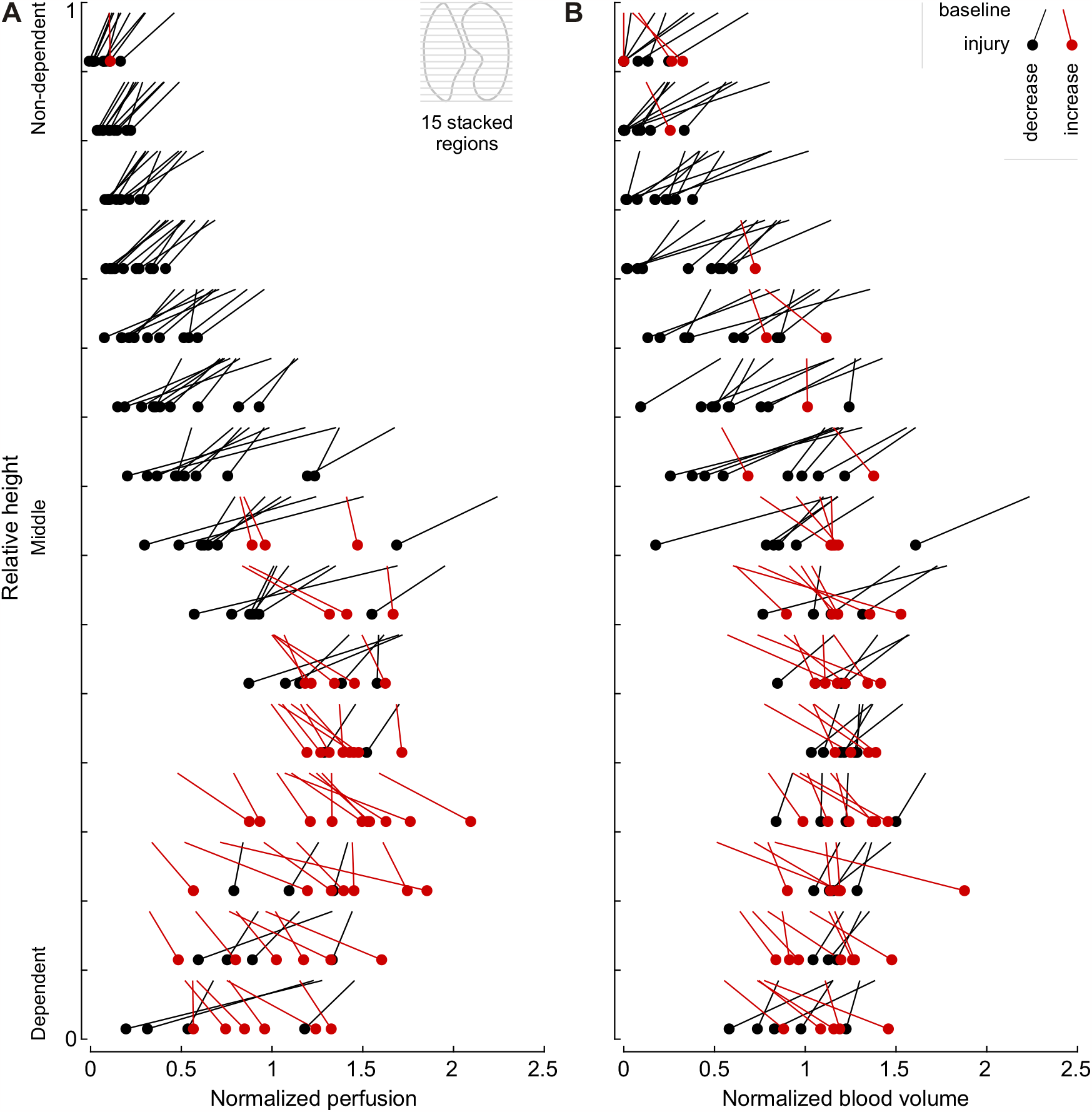
Vertical profiles of perfusion (A) and blood volume (B) with height expressed relative to the lung mask. Changes between baseline and lung injury are visualized as comets within each layer. The profiles and the changes are consistent with the example images in Fig. 1. Note the magnitude and frequency of decreases in perfusion and blood volume in non-dependent regions in contrast to the increases in the dependent half of each profile.

### Relationship of regional changes in adjusted Q_n_ and F_bn_ referenced to baseline

Although for the most part, changes in the adjusted Q_n_ and F_bn_ between baseline and injury were similar in direction, the magnitude of these changes for each variable were different and changed according to the vertical gradient (Fig. 4A and 4B). In the most non-dependent region, decreases in both Q_a_ and F_ba_ had lower magnitudes than in other regions because they were constrained by their low values at baseline. However, the longitudinal decrease to very low values of both F_ba_ and Q_a_ demonstrates the very high risk of capillary collapse in the most non-dependent regions in early ARDS with endotoxin exposure. An important insight from comparing regional changes in Q_a_ and F_ba_ visualized as slopes is the substantial variability in the regional responses rather than a common trend in their longitudinal changes (Fig. 4B).

**Figure 4.**
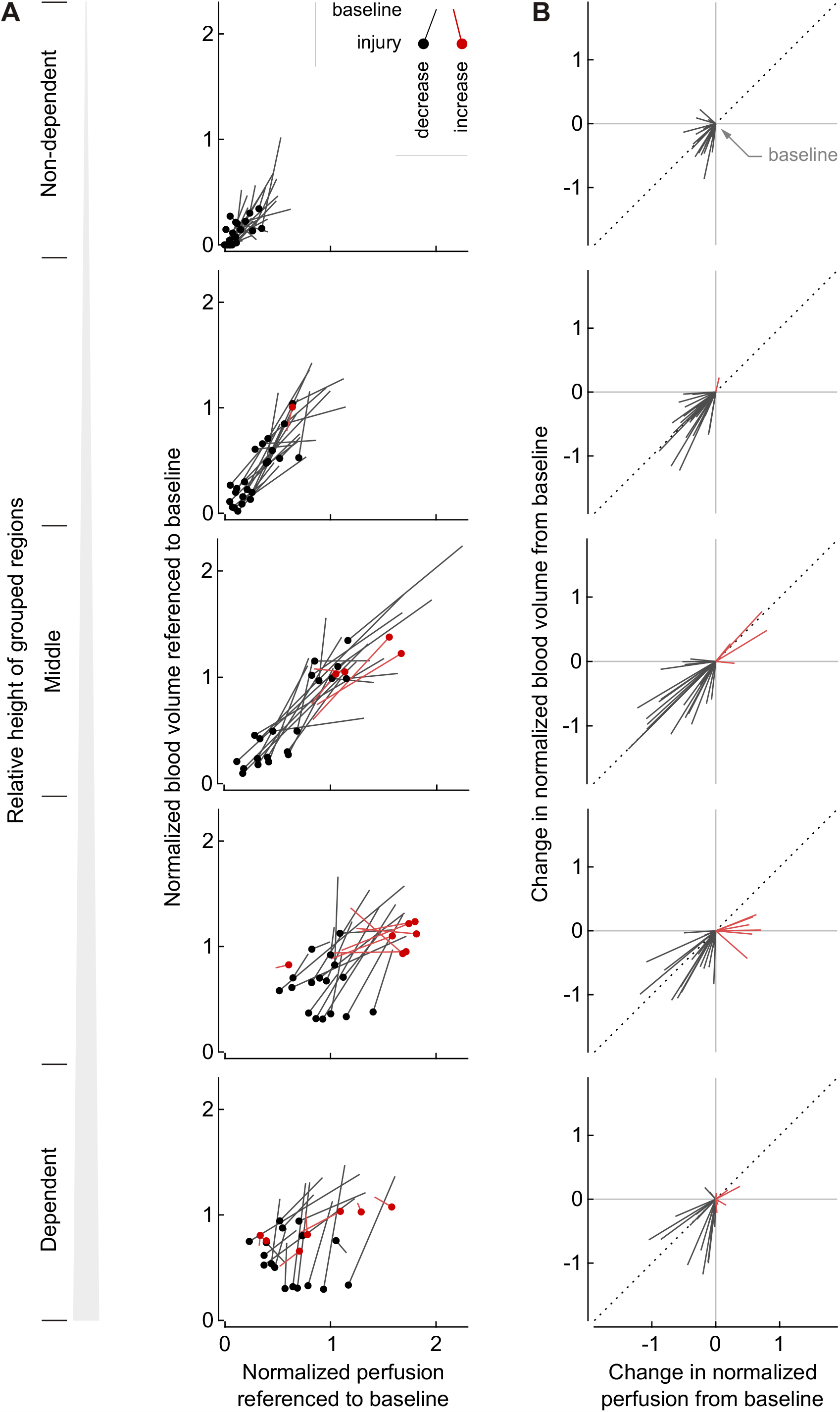
Longitudinal changes in normalized perfusion and blood volume between baseline and injury grouped at five levels of relative height show differences in regional responses (A), and their changes starting from a common baseline point visualize deviations in Q-Fb relationships (B). The longitudinal changes in normalized perfusion and blood volume are referenced to baseline values so that changes in cardiac output and pulmonary blood volume inside the lungs in the field of view are taken into account. Note the decrease in perfusion and blood volume in non-dependent regions dropping from low to very low or zero (A), which constrains the magnitude of the changes from baseline (B) and indicates a shift towards regional alveolar dead space. Deviations from the 45-degree slope (dotted line) show that the longitudinal changes in normalized blood volume were not equal to the changes in normalized perfusion.

## Discussion

Using advanced image analysis in an experimental model of lung injury representative of early ARDS with endotoxin exposure, we found that: 1) lung injury affects the relationship between normalized regional blood volume F_bn_ and normalized perfusion Q_n_, 2) the relationship changed from linear at baseline to curvilinear during lung injury, and 3) changes in the relationship were associated with differences in regional changes in Q_n_ compared to changes in F_bn_.

These findings are important mainly for the following reasons: 1) They challenge the use of blood volume as a quantitatively accurate surrogate for regional pulmonary perfusion. 2) They suggest that the blood volume-perfusion relationship, rather than being passive like a physical equation, is affected by vasoactive blood flow regulation, including HPV. 3) They demonstrate regional differentiation in the responses to endotoxin, including relative regional decreases in resistance in dependent regions blunting HPV in contrast to increases in the global PVR, and the risk for a capillary collapse in non-dependent regions.

In human and animal studies, investigators have previously found good correlations between the regional blood volume and perfusion, not normalized by regional differences in tissue fraction, using DECT (5,6) or PET (21,22) for blood volume compared to a perfusion imaging method. These correlations suggested a linear relationship between the two parameters, which is consistent with the relationship identified in our study at baseline. However, it has not been investigated if lung injury during long-term (20h) mechanical ventilation with endotoxin exposure may affect this relationship, which has consequences in clinical practice for longitudinal follow-up of changes in patients’ lung perfusion. Additionally, a PET study in healthy nonsmoking subjects has shown a curvilinear rather than a linear relationship (12).

Regional blood volume imaging relies on perfusion for tracer transport into the blood pool. For longitudinal quantitative assessments of regional changes in perfusion based on blood volume imaging, the relationship between blood volume and perfusion must remain the same when the patient’s lung conditions change. However, regional vasoactive blood flow regulation may not have the same effect on blood volume and perfusion. Additionally, changes in both total perfusion and regional vascular conditions interact with the structure and function of the pulmonary vascular tree such as regional differences in vascular properties, parenchymal tethering, blood clots and local vascular obstructions. We found a significant change in the relationship between blood volume and perfusion between baseline and the lung injury representative of early ARDS in an acute animal model.

We speculate that the higher dispersion of residuals relative to the modelled F_bn_- Q_n_ relationship at baseline compared to lung injury (Fig. 2) could be related to variations in the vasoactive blood flow regulation in the initially healthy lungs under conditions of anaesthesia that are altered by the effects of endotoxin exposure leading to both increased PVR and blunting of HPV(23). Interestingly, the quadratic F_bn_-Q_n_ relationship during endotoxin exposure is, despite the vasoactive effects of endotoxin, consistent with the behaviour expected of a passive vascular tree. The physical equation for laminar flow suggests that vascular conductance determining the flow has a quadratic relationship with the cross-sectional area of the vessel, and this area is proportional to blood volume if the length of the vessel is equal. A similar curvilinear relationship between perfusion and blood volume has also been found in a PET study in healthy subjects (12). A plausible explanation for the change in the F_bn_-Q_n_ relationship from linear at baseline and to curvilinear at lung injury could be that endotoxin exposure causes significant regional vasoactive rather than passive responses.

Additionally, higher variations in the normal vasoactive blood flow regulation at baseline may have increased the residuals during model comparisons to a degree that the BIC of a quadratic model, similar to the relationship for lung injury, lost its advantage over the linear model, which may have contributed to the selection of the linear model for baseline in contrast to the curvilinear model for lung injury representative of early ARDS (Table 2A and 2B, and online supplement Table 1A).

The distributions of Q_n_ and F_bn_ over height (Fig. 3) show that regions with low perfusion and low blood volume were located in nondependent areas. Also, Q_n_ and F_bn_ further decreased in most cases after lung injury, which could lead to capillary collapse. Contributing factors could be the vasoconstrictive effect of endotoxin, increased plateau pressures decreasing the intracapillary-to-alveolar pressure difference, and ‘flow stealing’ as perfusion shifts to more dependent regions.

Limitations of our study include that we present data from a small sample size which could make it harder to reach a statistical significance. However, we show very consistent characteristics among animals not only at baseline and during lung injury but also for the change from baseline to lung injury. We studied endotoxin infusion as a model of early ARDS which does not represent the entire clinical spectrum of this syndrome. Also, our study is limited to changes during lung injury representative of early ARDS (20 hours model) and the relevance or long-term behaviour of our findings are unknown. Nevertheless, the studied period was sufficient to capture changes in F_bn_ and Q_n_. Finally, the size of ROIs in our study may be larger than in other studies. However, larger ROIs have the advantage that they reduce the random measurement errors by averaging over a larger volume compared to smaller ROIs.

## Conclusions

In an experimental study, lung injury representative of early ARDS changed the relationship between regional blood volume and perfusion from linear to curvilinear. Effects of endotoxin exposure on the vasoactive blood flow regulation were most likely the key factor for this change, suggesting a primarily vasoactive rather than passive response and limiting the quantitative accuracy of blood volume imaging as a surrogate for regional perfusion and the interpretation of longitudinal changes.

## Acknowledgements

The authors thank Steve Weise^1^ for the expert support with PET imaging and the cyclotron staff John A. Correia^1^, Ph.D., and David F. Lee^1^, B.S. for the preparation of the radioisotopes. ^1^Department of Radiology (Nuclear Medicine and Molecular Imaging), Massachusetts General Hospital, Boston, MA, USA

## Authors contribution

AS and TW conceived the hypothesis and designed the analysis. GCMR contributed to the data analysis. MFVM designed the original experiments, and MFVM, MT, NDP, TJW, and TW performed the animal experiments. AS drafted the manuscript that all authors reviewed, edited, and approved.

## Data availability

Data are available upon reasonable request.

## Competing interest

Authors declare that they have no competing interests.

## Notes

### Competing Interest Statement

The authors have declared no competing interest.

