## Supplemental data for "Regional pulmonary perfusion, blood volume, and their relationship change during early ARDS in an experimental study"

4. Biomedical Engineering Program, Universidade Federal do Rio de Janeiro, Rio de Janeiro, Brazil.

5. Medical Intensive Care Unit, Hôpital Henri Mondor, Assistance Publique - Hôpitaux de Paris, Créteil, France

6. Divisao de Pneumologia, Instituto do Coracao, Hospital Das Clinicas HCFMUSP, Faculdade de Medicina, Universidade de Sao Paulo, SP, Brazil

7. Ultrasound Digital Solutions, GE Healthcare, San Mateo, California, USA.

8. Department of Anesthesiology, Columbia University Irving Medical Center,New York, New York, USA.



Funding: NIH-NHLBI grant R01-HL121228. GCMR was also funded by CAPES, Ministério da Educação do Brasil (scholarship 6344/15-1).



**Corresponding author**

Tilo Winkler, Department of Anesthesia, Critical Care and Pain Medicine, Massachusetts General Hospital and Harvard Medical School, Boston, Mass, USA.
ORCID: https://orcid.org/0000-0002-7276-5550

### Methods

*Subjects and Experimental design*

The experimental design has been previously described in detail (1). Briefly, sheep were anesthetized, intubated and mechanically ventilated. Anaesthesia was maintained with a continuous infusion of ketamine and propofol titrated to heart rate and blood pressure. Paralysis was established with a bolus of pancuronium (0.1 mg/kg) at induction and repeated every 90 min (0.02-0.04 mg/kg). Once an adequate sedation level was reached, the animals were intubated and connected to a Servo Ventilator 900C (Maquet, Solna, Sweden) in volume control, PEEP 5cm$H_2O$, tidal volume 6ml/kg, inspired $O_2$ fraction to maintain an arterial oxygen saturation ≥ 90% and respiratory rate to keep a $PaCO_2$ between 32 and 45 mmHg. A Swan Ganz catheter was introduced trough the right jugular vein for monitoring pulmonary artery pressure, mixed venous blood samples extraction and cardiac output measurement. An arterial line was canalized at the right femoral artery for arterial pressure monitoring and blood samples extraction. Volumetric capnography and respiratory mechanics were evaluated by means of a Nico monitor (Philips, Wallingford, CT).

Once animal instrumentation was completed, baseline (BL) measurements were performed, which consisted of arterial and mixed venous blood gases, systemic and pulmonary hemodynamics data collection, thermodilution cardiac output and a set of dynamic $^{13}$NN and $^{18}$F-FDG PET image acquisitions.

*PET images acquisition*

The PET imaging equipment, protocol, and processing methods have been previously presented in detail (Musch et al. 2007) (Musch et al. 2008) (O'Neill et al. 2003) (Vidal Melo et al. 2003). Briefly, we collected 15 PET transverse slices of 6.5-mm thickness, which provide three-dimensional information over a 9.7-cm-long field of view, estimated to encompass approximately 70% of the total sheep lung volume. Resulting images consisted of an interpolated matrix of 128 × 128 × 15 voxels with a size of 2.0 × 2.0 × 6.5 mm each. Three different types of PET images were acquired.

1.Transmission scans were obtained at baseline (0 h) and at 20 h using a rotating pin source of $^{68}$Ge for 10 min. Transmission scans were used for the attenuation correction of the corresponding emission scans, to demarcate the lung field, and to determine the fraction of gas ($F_{gas}^*$), which was computed by scaling the transmission scan to a range of 0 (defined as the attenuation of heart) to 1 (defined as the attenuation of air). Fgas of a region of interest (ROI) was calculated as the mean value of $F_{gas}^*$ within the ROI.

2.$^{13}$NN (nitrogen) emission scans were obtained at baseline and at the end of the 20-h mechanical ventilation period for the assessment of regional ventilation, perfusion, and shunt as previously reported with the $^{13}$NN-saline method. The imaging protocol was started with a tracer-free lung. The ventilator was stopped at the end of inhalation, and the airway pressure was equilibrated to a value equal to the mean airway pressure during ventilation. Then a 30- to 40-ml bolus of $^{13}$NN-saline solution was then centrally injected at a rate of 10 ml/s. Simultaneously, the collection of the dynamic PET scan with a sequence of frames (8 × 2.5, 4 × 10, 6 × 10, and 4 × 30 s) was started. After an apnea period of 60 s including the first 12 frames of the PET sequence, mechanical ventilation was restarted so that the washout phase consisted of 10 frames. The total imaging sequence lasted 4 min. Images of regional perfusion $Q_r^*$ were obtained by calculating the $^{13}$NN activity reached at the end of the breathhold plus the shunt fraction equal to the relative height of a peak prior to the plateau, if there is shunt. $Q_r^*$ of each ROI is equal to the average of the voxels within the ROI.

3. $^{18}$F-FDG emission scans: after $^{13}$NN clearance, $^{18}$F-FDG dissolved in 8 ml saline (approximately 40 MBq at 0 h and 200 MBq at 6 and 20 h) was infused at a constant rate through the jugular catheter for 60 s. Starting at the beginning of $^{18}$F-FDG infusion, sequential PET frames (9 × 10, 4 × 15, 1 × 30, 7 × 60, 15 × 120, 1 × 300, and 3 × 600 s) were acquired for 75 min. Pulmonary arterial blood was sampled at 5.5, 9.5, 25, 37, and 42.5 min, and plasma activity was measured in a well counter in order to calibrate an image-derived input function (2) for the three-compartment model (3). Regional $^{18}$F-FDG kinetics were fitted to a three-compartment model consisting of an intravascular compartment, a tissue compartment representing the concentration of $^{18}$F-FDG available for phosphorylation (i.e., $^{18}$F-FDG that is a substrate for hexokinase), and a metabolized compartment accounting for the concentration of $^{18}$F-FDG that has been phosphorylated by hexokinase (3,4). In this analysis, the $^{18}$F-FDG net uptake rate (Ki), a measure of cellular metabolic activity, is expressed as Ki = Fe·k3, where k3 is the rate of $^{18}$F-FDG phosphorylation and Fe is the fractional distribution volume of $^{18}$F-FDG that is in the tissue but not phosphorylated. In addition, kinetic analysis provides constants k1 (transfer rate from plasma to tissue), k2 (transfer rate from tissue to plasma), and $F_B$ (the blood volume fraction in the ROI).

### Statistical Analysis

The primary objective of this study was to determine if the relationship between Fb and Q during early ARDS is different from baseline. To answer it, we performed a kernel density estimator (KDE) test under the null hypothesis that the Fb vs Q data at early ARDS are from the same distribution as the baseline data. As secondary analysis aiming to identify models that describe the relationship between Fb and Q, we compared six different relationships between Fb and Q at baseline and early ARDS conditions and the assignment of Fb and Q to the independent and dependent parameter using the Bayesian information criterion (BIC) as primary parameter for the identifying the best model. The six relationships included zero to second order polynomial and exponential functions. Secondary parameters for checking the plausibility of the model identification were Akaike's information criterion (AIC) and root of mean squared errors. After performing this analysis we constructed polar plots of the changes in Fba vs Qa pairs from baseline to ARDS trying to describe a regional distribution of the model effect on these variables. Despite of several measures were taken at each animal, no repeated measures analysis was performed because we expect that the between animals differences were majorly removed by the mean normalization. A p value < 0.05 was assumed as significant. Values are expressed as mean ± standard deviation unless otherwise specified. The effect of the early ARDS model in physiologic variables was evaluated by applying a paired t test.  The computational analysis was performed using Matlab (Mathworks, Natick, MA). The statistical analysis was performed using Stata (v14.2; StataCorp LLC, College Station, TX), the R Statistical Software (v4.2.1; R Core Team 2022), and the ks package (v1.13.5) (5) for the KDE test.

### Extended Discussion

In principle, blood flow requires an intravascular blood volume so that a relationship between regional perfusion and blood volume could be expected. However, punctuated vascular obstructions could result in regional blood volume with very low blood flow, and cause significant deviations from a relationship between blood volume and flow under normal conditions without such obstructions. Additionally, lung injury and other conditions can affect vascular smooth muscle tone regulating regional blood flow. It is known that regional pulmonary perfusion and blood volume are related, but not if they have a fixed relationship or if the parameters can have different relevant information, and reflect different aspects of vascular function, which is relevant for the understanding of the pathophysiology of lung injury during early ARDS and the interpretation of DECT images.

Supplementary tables

Table 1

A - Baseline

| Model | Root MSE | AIC |
|---|---|---|
| $y = ax$ | 0.294 | 59.370 |
| $y = a + bx$ | 0.255 | 17.956 |
| $y = ax + bx^2$ | 0.258 | 21.575 |
| $y = a + bx + cx^2$ | 0.253 | 16.199 |
| $y = e^{ax}$ | 0.402 | 153.455 |
| $y = ae^{bx}$ | 0.268 | 32.403 |

B – Lung injury

| Model | Root MSE | AIC |
|---|---|---|
| $y = ax$ | 0.247 | 7.433 |
| $y = a + bx$ | 0.213 | -36.605 |
| $y = ax + bx^2$ | 0.165 | -112.034 |
| $y = a + bx + cx^2$ | 0.166 | -110.054 |
| $y = e^{ax}$ | 0.514 | 225.316 |
| $y = ae^{bx}$ | 0.268 | 32.894 |

Table 2

A – Baseline

| Dependent | Independent | Model | a | b | c | BIC |
|---|---|---|---|---|---|---|
| $F_{bn}$ | $Q_n$ | $y = ax$ | 0.980 | | | 62.381 |
| $Q_n$ | $F_{bn}$ | | 0.934 | | | 55.275 |
| $F_{bn}$ | $Q_n$ | $y = a + bx$ | 0.320 | 0.690 | | 23.977 |
| $Q_n$ | $F_{bn}$ | | 0.071 | 0.870 | | 58.828 |
| $F_{bn}$ | $Q_n$ | $y = ax + bx^2$ | 1.433 | -0.357 | | 27.596 |
| $Q_n$ | $F_{bn}$ | | 0.966 | -0.026 | | 60.130 |
| $F_{bn}$ | $Q_n$ | $y = a + bx + cx^2$ | 0.204 | 1.008 | -0.168 | 25.231 |
| $Q_n$ | $F_{bn}$ | | 0.124 | 0.723 | 0.081 | 63.206 |
| $F_{bn}$ | $Q_n$ | $y = e^{ax}$ | 0.078 | | | 156.465 |
| $Q_n$ | $F_{bn}$ | | 0.024 | | | 202.305 |
| $F_{bn}$ | $Q_n$ | $y = ae^{bx}$ | 0.503 | 0.650 | | 38.425 |
| $Q_n$ | $F_{bn}$ | | 0.367 | 0.881 | | 69.412 |

B – Lung Injury

| Dependent | Independent | Model | a | b | c | BIC |
|---|---|---|---|---|---|---|
| $F_{bn}$ | $Q_n$ | $y = ax$ | 0.967 | | | 10.437 |
| $Q_n$ | $F_{bn}$ | | 0.961 | | | 9.590 |
| $F_{bn}$ | $Q_n$ | $y = a + bx$ | 0.214 | 0.780 | | -30.598 |
| $Q_n$ | $F_{bn}$ | | -0.075 | 1.031 | | 10.844 |
| $F_{bn}$ | $Q_n$ | $y = ax + bx^2$ | 1.684 | -0.538 | | -106.026 |
| $Q_n$ | $F_{bn}$ | | 0.502 | 0.390 | | -10.077 |
| $F_{bn}$ | $Q_n$ | $y = a + bx + cx^2$ | 0.004 | 1.674 | -0.532 | -101.042 |
| $Q_n$ | $F_{bn}$ | | 0.055 | 0.370 | 0.459 | -6.610 |
| $F_{bn}$ | $Q_n$ | $y = e^{ax}$ | 0.097 | | | 228.320 |
| $Q_n$ | $F_{bn}$ | | 0.040 | | | 278.075 |
| $F_{bn}$ | $Q_n$ | $y = ae^{bx}$ | 0.415 | 0.775 | | 38.902 |
| $Q_n$ | $F_{bn}$ | | 0.212 | 1.375 | | 22.047 |

3. Sokoloff L, Reivich M, Kennedy C, Des Rosiers MH, Patlak CS, Pettigrew KD, et al. The [14C]deoxyglucose method for the measurement of local cerebral glucose

utilization: theory, procedure, and normal values in the conscious and anesthetized albino rat. J Neurochem. 1977 May;28(5):897–916.

4. Dittrich AS, Winkler T, Wellman T, de Prost N, Musch G, Harris RS, et al. Modeling 18F-FDG kinetics during acute lung injury: experimental data and estimation errors. PLoS One. 2012;7(10):e47588.

5. Duong [aut T, cre, Wand M, Chacon J, Gramacki A. ks: Kernel Smoothing [Internet]. 2022 [cited 2022 Aug 22]. Available from: https://CRAN.R-project.org/package=ks
